# The ribosome-associated quality control factor TCF25 imposes K48 specificity on Listerin-mediated ubiquitination of nascent chains by binding and specifically orienting the acceptor ubiquitin

**DOI:** 10.1101/2024.10.17.618946

**Authors:** Irina S. Abaeva, Alexander G. Bulakhov, Christopher U. T. Hellen, Tatyana V. Pestova

## Abstract

Polypeptides arising from interrupted translation undergo proteasomal degradation by the ribosome-associated quality control (RQC) pathway. The ASC-1 complex splits stalled ribosomes into 40S subunits and nascent chain-tRNA-associated 60S subunits (60S RNCs). 60S RNCs associate with NEMF that promotes recruitment of the RING-type E3 ubiquitin (Ub) ligase Listerin (Ltn1 in yeast), which ubiquitinates nascent chains. RING-type E3s mediate the transfer of Ub directly from the E2∼Ub conjugate, implying that the specificity of Ub linkage is determined by the given E2. Listerin is most efficient when it is paired with promiscuous Ube2D E2s. We previously found that TCF25 (Rqc1 in yeast) can impose K48- specificity on Listerin paired with Ube2D E2s. To determine the mechanism of TCF25’s action, we combined functional biochemical studies and AlphaFold3 modeling and now report that TCF25 specifically interacts with the RING domain of Listerin and the acceptor ubiquitin (Ub^A^) and imposes K48-specificity by orienting Ub^A^ such that its K48 is directly positioned to attack the thioester bond of the Ube2D1∼Ub conjugate. We also found that TCF25 itself undergoes K48-specific ubiquitination by Listerin suggesting a mechanism for the reported upregulation of Rqc1 in the absence of Ltn1 and the observed degradation of TCF25 by the proteasome *in vivo*.

## INTRODUCTION

To ensure the accuracy of gene expression and to limit the accumulation of incomplete, potentially cytotoxic polypeptides that arise from ribosomal stalling during eukaryotic protein synthesis, eukaryotes employ highly conserved mRNA and protein quality control systems. The No-Go decay surveillance mechanism targets mRNAs on which ribosomes are stalled by e.g. stable secondary structures, rare codons, polyA stretches or damaged RNA bases (d’Orazio and Green, 2021; Yip and Shao, 2021) whereas corresponding nascent chain (NC) polypeptides arising from interrupted translation are degraded by ribosome-associated quality control (RQC) pathways via the ubiquitin-proteasome system (Joazeiro, 2019; Sitron and Brandman, 2020; Inada and Beckmann, 2024). Their physiological importance is underlined by the growing number of pathological states linked to defects in the RQC process (Lu, 2022; McGirr et al., 2024).

In RQC, stalled ribosomes K63-polyubiquitinated by the E3 ubiquitin ligase ZNF598 (Hel2 in yeast) (Juszkiewicz and Hegde, 2017; Juszkiewicz et al., 2018; Matsuo et al., 2017; Sundaramoorthy et al., 2017) are dissociated by the ASC-1 (activating signal cointegrator 1) complex (the RQC-Trigger complex in yeast) into 40S subunits and NC-tRNA/60S ribosome-nascent chain complexes (60S RNCs) (Hashimoto et al., 2020; Juszkiewicz et al., 2020; Matsuo et al., 2020; Miścicka et al., 2024). This exposes the P site tRNA and the interface surface on the 60S subunit, allowing binding of NEMF (Rqc2 in yeast), which recruits to the 60S RNCs the RING-type E3 ubiquitin ligase Listerin (Ltn1 in yeast) (Defenouillère et al., 2013; Shao et al., 2015; Shen et al., 2015). Listerin’s N-terminal domain binds to NEMF/Rqc2 and the sarcin-ricin loop of the 60S subunit and is linked by a long HEAT repeat-containing region to the C-terminal RWD and RING domains, which bind near the nascent polypeptide exit tunnel that allows ubiquitination of the protruding NCs within a small window outside the exit tunnel (Lyumkis et al., 2014; Doamkepor et al., 2016; Kostova et al., 2017; Shao et al., 2015; Shen et al., 2015; Tesina et al., 2023). TCF25 (Rqc1 in yeast) stimulates ubiquitination and importantly, imposes K48 specificity on it (Kuroha et al., 2018). The oligo-Ub tag on Ub-NCs recruits the AAA ATPase VCP/p97 (cdc48 in yeast) with its heterodimeric Npl4/Ufd1 cofactor (Brandman et al., 2012; Verma et al., 2013; Defenouillère et al., 2013), which extracts proteasome-degradable Ub-NCs linked to three 3’-terminal tRNA nucleotides from the exit tunnel after specific cleavage in the tRNA acceptor arm by ANKZF1 (Vms1 in yeast) (Kuroha et al., 2018; Verma et al., 2018; Yip et al., 2019). The K48 specificity of Ub linkage enforced by TCF25 renders the ubiquitinated NC an optimal substrate for binding by cdc48/Ufd1/Npl4 (Tsuchiya et al., 2017; Sato et al., 2019; Williams et al., 2023) and subsequent degradation by the proteasome (Thrower et al., 2000). However, the mechanism by which TCF25 performs its function remained unknown.

Protein ubiquitination begins with ATP-dependent activation of Ub by thioester linking of its C-terminus to the active site Cys of ubiquitin-activating enzyme E1. Ub is then transthiolated to the Cys residue of a catalytic ubiquitin conjugating (UBC) domain of ubiquitin-conjugating enzyme E2, yielding an E2∼Ub conjugate. E2-conjugated Ub is designated as "donor" Ub (Ub^D^). Selection of a protein substrate is determined by E3 ubiquitin ligases that bind and bring together both E2∼Ub^D^ and a target. Ubiquitination of a substrate occurs by a nucleophilic attack of the amino group of a Lys (or less commonly, of an N-terminal) residue on the E2∼Ub^D^ thioester linkage, forming a peptide bond with the C-terminus of Ub. The attached Ub molecule can now attack other E2∼Ub^D^ conjugates forming Ub chains. Ub that attacks the conjugate is designated as acceptor Ub (Ub^A^). E3 ligases function by two different mechanisms. HECT-type E3 ligases form a catalytic intermediate with Ub^D^ (also by transthiolation) before its transfer to the substrate and therefore can determine the specificity of Ub linkage (Lorenz, 2018). In contrast, RING-type E3 ligases mediate the transfer of ubiquitin directly from E2∼Ub to the substrate, in which case the specificity of linkage is determined by the given E2 (Stewart et al., 2016).

The ∼50-70 aa-long RING domains have the consensus sequence C1-X_(2)-_C2-X_(9–39)-_C3-X_(1–3)-_H4- X_(2–3)-_C/H5-X_(2)-_C6-X_(4–48)_C7-X_(2)_C8, which forms two Zn^2+^-coordination sites with a unique “cross-brace” arrangement: Cys1/Cys2 and Cys(His)5/Cys6 bind the first and Cys3/His4 and Cys7/Cys8 bind the second Zn^2+^ ion (Metzger et al., 2014). In addition to their scaffolding function of bringing a substrate and E2∼Ub together, RING domains promote a closed, active conformation of the E2-Ub^D^ conjugate in which the flexibly linked Ub is constrained by an interaction with a E2 surface close to its active sites rendering the thioester bond susceptible to nucleophilic attack (Lorick et al., 1999; Buetow and Huang, 2016; Branigan et al., 2020). The RING domain binds the E2 on a surface distant from the active site Cys, allosterically activating a closed conformation of E2-Ub^D^. An ‘allosteric linchpin’ residue (usually Arg), which is part of a conserved ϕ-x-K/R motif in which x is Cys8 of the RING domain and ϕ is a hydrophobic residue, engages both Ub and E2 through hydrogen bonding (Dou et al., 2012; Plechanovová et al., 2012; Pruneda et al., 2012). In addition, RING-type E3s also promote the closed conformation of E2-Ub through diverse direct interactions with Ub^D^.

Listerin is a RING-type E3 ligase, implying that the specificity of its linkage will be determined by the paired E2. Although K48 linkage would make the ubiquitinated NC an optimal substrate for proteasomal degradation, Listerin is most active with Ube2D and Ube2E family E2s (Shao et al., 2013; Kuroha et al., 2018), which are both known for their promiscuity, implying that they bind acceptor Ub (Ub^A^) at different sites and/or orientations and form all linkages (Jin et al., 2008; Stewart et al., 2016). We have previously found that TCF25 imposes K48 specificity on ubiquitination by Listerin in combination with Ube2D or Ube2E E2s (Kuroha et al., 2018), but the mechanism of its action remains unknown. Another example of a separate protein influencing E2 specificity is the K63-specific ubiquitination by Ube2N, which is determined by a non-catalytic E2-like co-factor, Ube2V1 or Ube2V2, that forms a heterodimer with the cognate E2, binds Ub^A^ and positions its K63 for attack on the E2∼UbD ester bond (Eddins et al., 2006; Branigan et al., 2015, 2020; Nakasone et al., 2022). The effect of TCF25 was Listerin-specific, and it did not influence the specificity of ubiquitination when Ube2D E2s were combined with other E3 ligases, e.g. Mdm2 (Kuroha et al., 2018), indicating that TCF25 is unlikely to perform its function through direct interaction with E2 enzymes.

Human TCF25 (also known as hNulp1) is 675 amino acids long (Olsson et al., 2002; Cai et al., 2006) and consists of a predominantly unstructured region that is predicted to contain two α-helices followed by a basic helix-loop-helix (bHLH) motif (aa 126-202) in which the basic region is unusually lysine-rich and the loop is unusually long relative to other bHLH proteins (Atchley and Fitch, 1997), a tetracopeptide α-helical DUF654 domain, and a weakly structured C-terminal region (Figure S1). TCF25 does not contain ubiquitin-binding domains, and its position in RQC complexes and interactions with ather RQC components are unknown. Here, we combined *in vitro* functional biochemical studies and AlphaFold3 modeling to determine the mechanism of TCF25 function.

## RESULTS

### The activity of TCF25 and other RQC components in promoting the active conformation of the E2∼Ub conjugate

In 60S RNCs, Listerin ubiquitinates the protruding NCs within a small window (Shao and Hegde, 2014; Shao et al., 2013, 2015; Kostova et al., 2017; Kuroha et al., 2018). To determine whether TCF25 is equally active over the entire Listerin-accessible window, we assayed its activity using sucrose density gradient (SDG)-purified 60S RNCs assembled in rabbit reticulocyte lysate on non-stop ϕ-VHP mRNAs comprising the ϕ-globin 5’ UTR, the coding region for the short structured villin headpiece and an unstructured part of Sec61ϕ ending with Val (Shao et al., 2013; Kuroha et al., 2018). Bearing in mind that the ribosomal exit tunnel accommodates 30-40 aa, the open reading frame included single Lys residues at positions 30 to 54 from the C-terminus. Ubiquitination was most efficient from Lys40 through to the last assayed position. TCF25 did not influence the ubiquitination window and was active along its entire length, stimulating ubiquitination by K48-only Ub and inhibiting ubiquitination by K48R Ub (Figure 1A).

**Figure 1.**
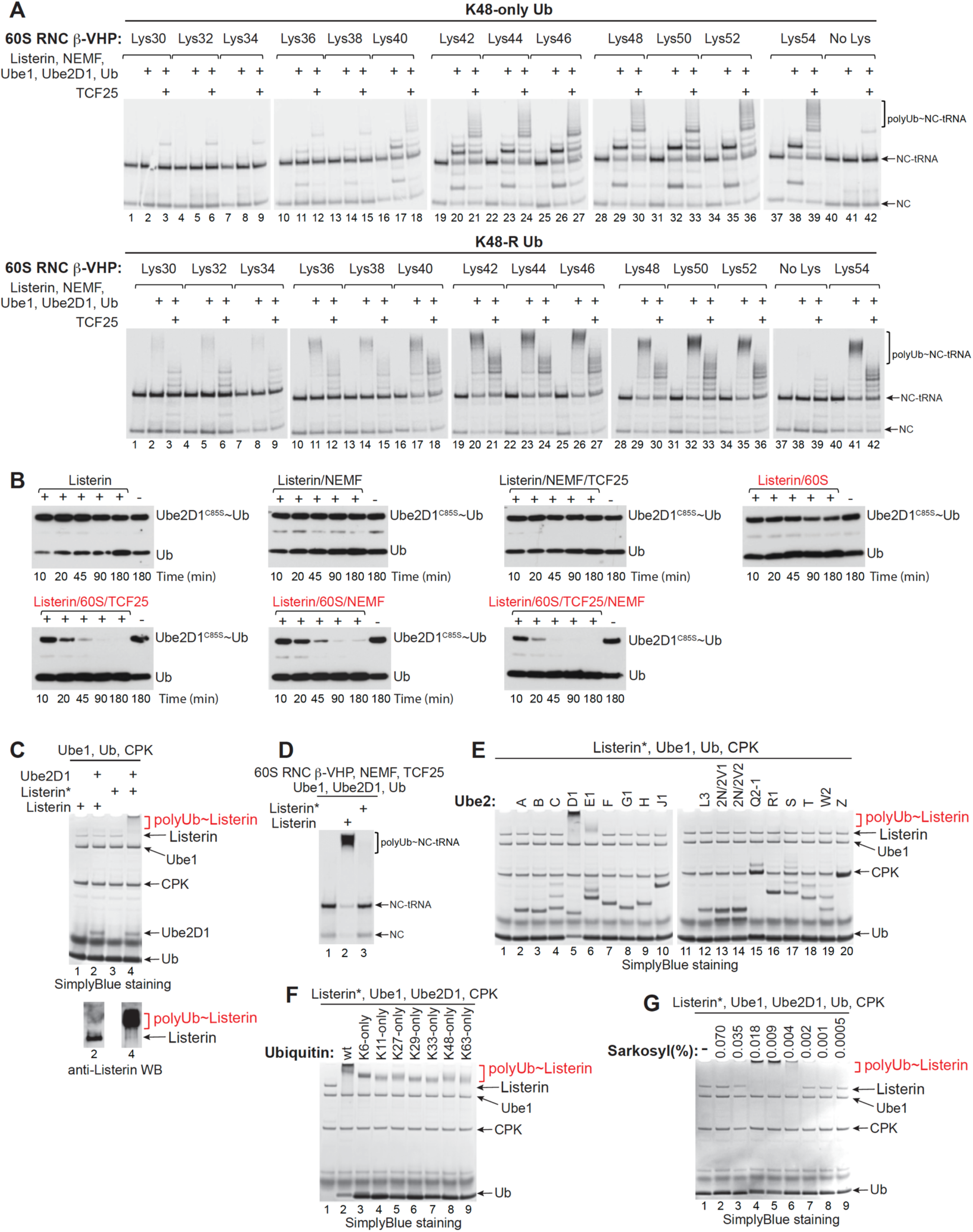
The activity of TCF25 along the ubiquitination window, Listerin-mediated hydrolysis of Ube2D1∼Ub conjugate and auto-ubiquitination of Listerin. (A) The activity of TCF25 along the ubiquitination window assayed by its influence on ubiquitination of ϕ-VHP-containing 60S RNCs with single Lys residues introduced at the indicated positions by Listerin/NEMF/Ube1/Ube2D1 with K48-only or K48R mutant Ub. Ubiquitination products were assayed by SDS-PAGE and visualized by Phosphoimager. (B) Time courses of Listerin-induced hydrolysis of Ube2D1^C85S^∼Ub conjugate in the presence of 60S subunits, TCF25 and NEMF, as indicated, assayed by western blotting. (C) Auto-ubiquitination of Listerin and Listerin* in the presence of Ube1/Ube2D1/*wt* Ub, assayed by SDS-PAGE followed by staining (upper panel) or western blotting (lower panel). (D) The activity of Listerin and Listerin* in ubiquitination of ϕ-VHP-containing 60S RNCs in the presence of NEMF/Ube1/Ube2D1/*wt* Ub, assayed by SDS-PAGE and visualized by Phosphoimager. (E) Auto-ubiquitination of Listerin* with Ube1, *wt* Ub and indicated Ube2 enzymes, assayed by SDS-PAGE followed by staining. (F) Auto-ubiquitination of Listerin* with Ube1, Ube2D1 and *wt* or indicated mutant Ub, assayed by SDS-PAGE followed by staining. (G) Auto-ubiquitination of Listerin with Ube1, Ube2D1 and *wt* Ub in the presence of Sarkosyl at indicated concentrations, assayed by SDS-PAGE followed by staining.

We next investigated whether in addition to imposing K48-specificity, TCF25 also stimulates formation of the active conformation of E2∼Ub. To evaluate the contribution of TCF25, NEMF and 60S subunits in Listerin-mediated promotion of the E2∼Ub active conformation, we assayed hydrolysis of the slowly hydrolysable Ube2D1^C85S^∼Ub conjugate linked by an oxyester rather than a thioester bond. No meaningful hydrolysis occurred without 60S subunits irrespective of the presence of NEMF and TCF25 (Figure 1B, upper row, three left panels). Slow, low-level hydrolysis was observed in the presence of 60S subunits alone (Figure 1B, upper right-hand panel). Inclusion of TCF25 or NEMF with 60S subunits strongly stimulated hydrolysis, and the most efficient hydrolysis of the conjugate occurred in the presence of 60S subunits and both NEMF and TCF25 (Figure 2B, lower panels). Regarding its ribosomal position and function during the quality control pathway, NEMF likely stimulated Ube2D1^C85S^∼Ub hydrolysis by promoting Listerin’s association with the 60S subunit, whereas stimulation by TCF25 could additionally involve a direct influence on the position/conformation of the RING domain and/or E2∼Ub.

**Figure 2.**
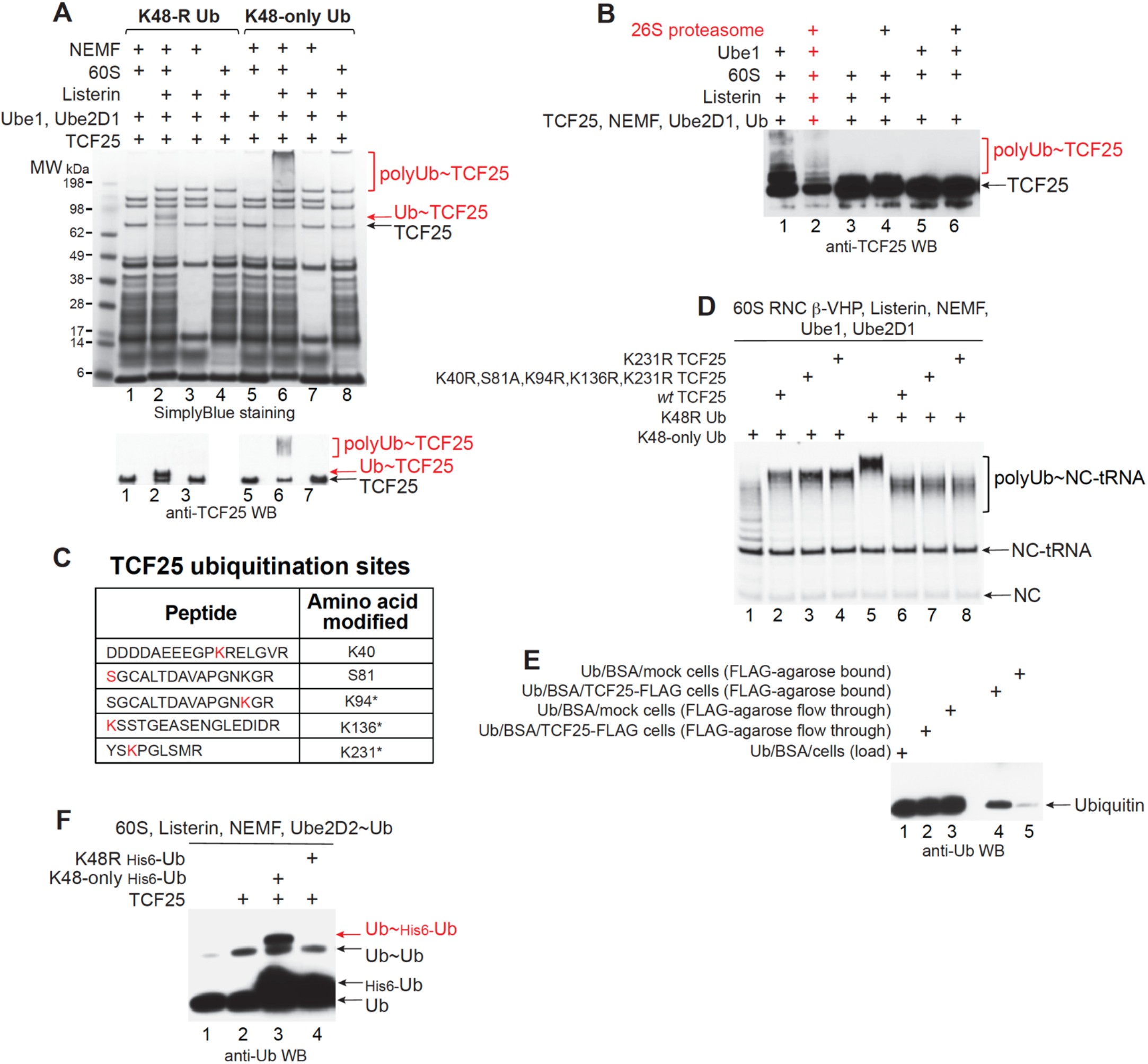
Listerin-mediated ubiquitination of TCF25 and TCF25-mediated formation of di-ubiquitin. (A) Listerin-mediated ubiquitination of TCF25 in the presence of 60S subunits, Listerin, Ube1, Ube2D1 and K48R or K48- only Ub, as indicated, assayed by SDS-PAGE followed by staining (upper panel) or by western blotting (lower panel). (B) Proteasomal degradation of TCF25 ubiquitinated by Listerin in the presence of other indicated components, assayed by western blotting. (C) TCF25 ubiquitination sites determined by mass spectrometry. * - indicates previously reported sites (https://www.phosphosite.org/). (D) The influence of *wt* and ubiquitination sites mutant TCF25 on ubiquitination of ϕ3-VHP- containing 60S RNCs by Listerin/NEMF/Ube1/Ube2D1 with K48-only or K48R mutant Ub, assayed by SDS-PAGE and visualized by Phosphoimager. (E) The interaction between purified recombinant ubiquitin and a FLAG-tagged TCF25- expressing Expi293 cell extract assayed by pull-down using anti-FLAG agarose beads followed by Western blotting. (F) TCF25-mediated formation of di-ubiquitin in reaction mixtures containing Listerin, 60S subunits, NEMF, Ube2D2∼Ub conjugate and His_6_-tagged K48-only or K48R Ub.

Interestingly, we occasionally purified Listerin (designated as Listerin*), which was not active in ubiquitination of nascent chains but, in contrast to active Listerin, underwent efficient auto-ubiquitination in the presence of Ube1, Ube2D1 and Ub (Figures 1C-D). Ubiquitination sites on Listerin* were dispersed throughout the protein, and many of them have been reported (https://www.phosphosite.org/) (Figure S2A). Listerin* retained canonical specificity to E2 enzymes, promoting promiscuous ubiquitination with Ube2D and Ube2E classes of E2s (Figures 1E-F). The presence of 60S subunits and other RQC components did not influence auto-ubiquitination of Listerin* (Figure S2B). The fact that native Listerin did not stimulate hydrolysis of E2∼Ub^D^ in the absence of 60S subunits and other RQC components indicates that individually it does not adopt the active state. Many E3 ubiquitin ligases exist in auto-inhibited conformations, in which regions outside catalytic domains interfere with their function. In some, auto-inhibition can be relieved by post-translational modification, e.g. phosphorylation (Gallagher et al., 2006; Vittal et al., 2015), but mass spec analysis did not reveal Listerin*-specific modifications, which suggested conformational changes due to local unfolding as the cause of auto-ubiquitination. Consistently, auto-ubiquitination of active Listerin was induced by unfolding when reaction mixtures were supplemented by the anionic detergent Sarkosyl at low concentrations of 0.004-0.018% (Figure 1G).

### Listerin-mediated ubiquitination of TCF25

Strikingly, we found that TCF25 itself also undergoes efficient ubiquitination in reaction mixtures containing Listerin, NEMF and 60S subunits. Importantly, TCF25 imposed K48-specificity on its own ubiquitination: in the presence of K48-only Ub, it was polyubiquitinated, whereas in the presence of K48R Ub, it was primarily mono-ubiquitinated (Figure 2A, lanes 2 and 6). NEMF was not strictly essential but stimulated the process. TCF25 polyubiquitinated with *wt* Ub was efficiently degraded by the 26S proteasome *in vitro* (Figure 2B), which suggests the mechanism for regulation of TCF25 levels *in vivo*. TCF25 was ubiquitinated at several sites (Figure 2C), some of which have been reported (https://www.phosphosite.org/). Arginine substitution of K231, the most frequently reported site of ubiquitination, or even of all ubiquitinated residues together did not influence the activity of TCF25 in ubiquitination of NCs in 60S RNCs (Figure 2D).

To test whether TCF25 can interact with Ub, we employed a FLAG pull-down assay using anti-FLAG agarose and a HEK293 cell extract expressing FLAG-tagged TCF25. TCF25 pulled down ubiquitin (Figure 2E) suggesting their direct interaction. This prompted us to assay the ability of TCF25 to promote formation of di-ubiquitin in reaction mixtures containing *wt* Ube2D2∼Ub conjugate, Listerin, NEMF, 60S subunits and His_6_-tagged K48-only or K48R Ub. Using His_6_-tagged mutant Ub allows Ub∼His_6_-Ub(mut) to be distinguished from *wt* Ub∼Ub, which can also form because the reaction mixtures would contain untagged *wt* Ub resulting from spontaneous and Listerin-induced hydrolysis of the *wt* Ube2D2∼Ub conjugate. In this system, TCF25 promoted efficient formation of di-ubiquitin from K48-only Ub but not from K48R Ub (Figure 2F).

Taken together, our data are consistent with a mechanism in which TCF25 functions by orienting acceptor ubiquitin (Ub^A^) such that K48 is positioned to attack the thioester bond of the E2∼Ub^D^ conjugate. In the case of K48R Ub, TCF25 would lock the complex in a non-productive state thereby inhibiting ubiquitination.

### AlphaFold3 model of the Listerin/TCF25/Ube2D1/Ub^D^/Ub^A^ complex and its validation

Our attempts to establish the position of TCF25 in *in vitro* assembled mammalian RQC complexes by cryo-electron microscopy were not successful, consistent with a previous report (Shao et al., 2015). Similarly, no density for Rqc1 was observed in cryo-EM structures of yeast RQC complexes even though they were purified by affinity to Rqc1 (Tesina et al., 2023). We therefore employed AlphaFold3 (Abramson et al., 2024) to generate a model for the interaction of TCF25, Listerin, Ube2D1 and two Ub molecules. In the predicted model, the structured core of TCF25 comprised by aa ∼175-655 interacts directly with the RING domain of Listerin and the Ub^A^, whereas the N-terminal (aa 1-174) and C-terminal (aa 656-676) regions are mostly unstructured and do not interact with other components of the complex (Figure 3). Strikingly, interaction with TCF25 orients Ub^A^ such that its K48 is positioned to attack directly the thioester bond of the E2∼Ub^D^ conjugate.

**Figure 3.**
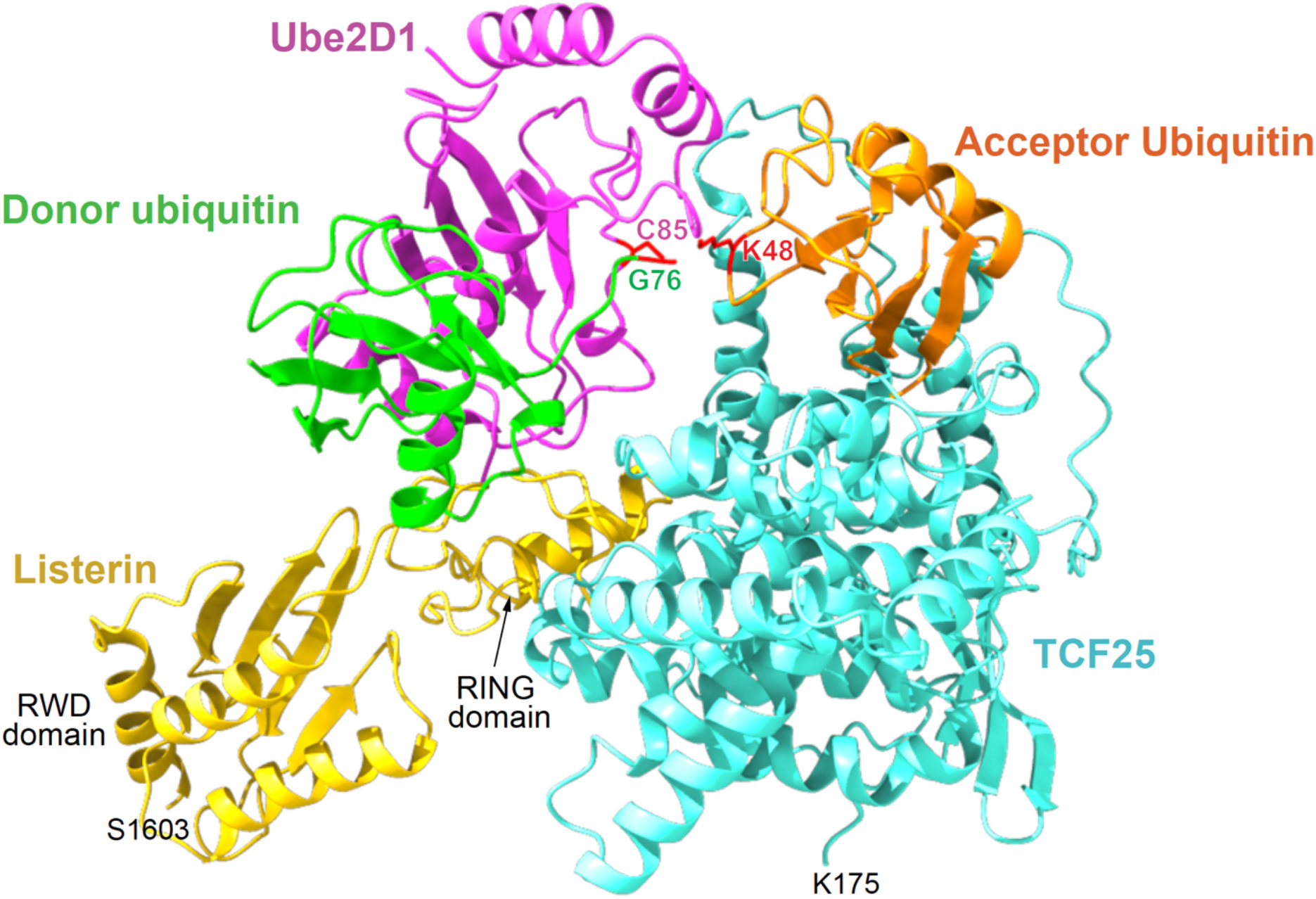
Model of the Listerin/TCF25/Ube2D1/Ub^D^/Ub^A^ complex. The AlphaFold3 model showing the complex between TCF25 (aa 175-655; light blue ribbon), the RING and preceding RWD domains of Listerin (aa 1603-1766; yellow ribbon), Ube2D1 (magenta ribbon), donor ubiquitin (green ribbon) and acceptor ubiquitin (orange ribbon). K48 in the acceptor ubiquitin, C85 in Ube2D1 and G76 in the donor ubiquitin are shown as red sticks. The unstructured N- and C- terminal regions of TCF25, as well as the N-terminal part of Listerin preceding the RWD domain which are not involved in the intermolecular interactions have been omitted for clarity.

To validate the AlphaFold3 model, we first assayed the predicted interaction between TCF25 and the RING domain. The strongest contacts seem to occur between several hydrophobic amino acids in α5-α6 helices of TCF25 (aa 188-202; Figure S1) and in the αA helix of the RING domain (aa 1743-1753; Figure S3A). Thus, the highly conserved F195 (Figure S1) in TCF25 can interact with L1746, F1766 and strongly with Y1747 and F1750 (Figure 4A). In addition to interaction with F195, Y1747 can contact L191 and I200, whereas F1766 may stack with Y194. Additionally, K1748 can interact with E203 in TCF25. Consistently, deletion of aa188-200 abrogated the activity of TCF25 in ubiquitinating NCs in 60S RNCs (Figure 4B, lanes 3 and 7). Abrogation of activity also resulted from the F195A substitution alone (Figure 4C, lanes 5 and 10). Ala substitution of the conserved E190 (Figure S1) had almost the same effect (Figure 4C, lanes 4 and 9), and even though E190 is not predicted to interact with Listerin, it is involved in multiple intramolecular contacts which are likely important for the proper positioning of the α5 helix (Figure 4A). The same abrogating effect on the activity of TCF25 was observed after deleting the preceding aa175-186 or by the E181A/R183A/L185A substitutions in this region (Figure 4D). Like E190, this region does not interact with the RING domain but could also contribute to proper positioning of the α5 helix through a network of intramolecular contacts (Figure 4A). Consistent with the loss of the activity in ubiquitination of nascent chains in 60S RNCs, ubiquitination of TCF25 mutants themselves was much less efficient than of wt TCF25 in reaction mixtures containing 60S subunits, Listerin and NEMF, and importantly, ubiquitination by K48R Ub was stronger than by K48-only Ub and yielded oligoubiquitinated proteins (Figure 4E).

**Figure 4.**
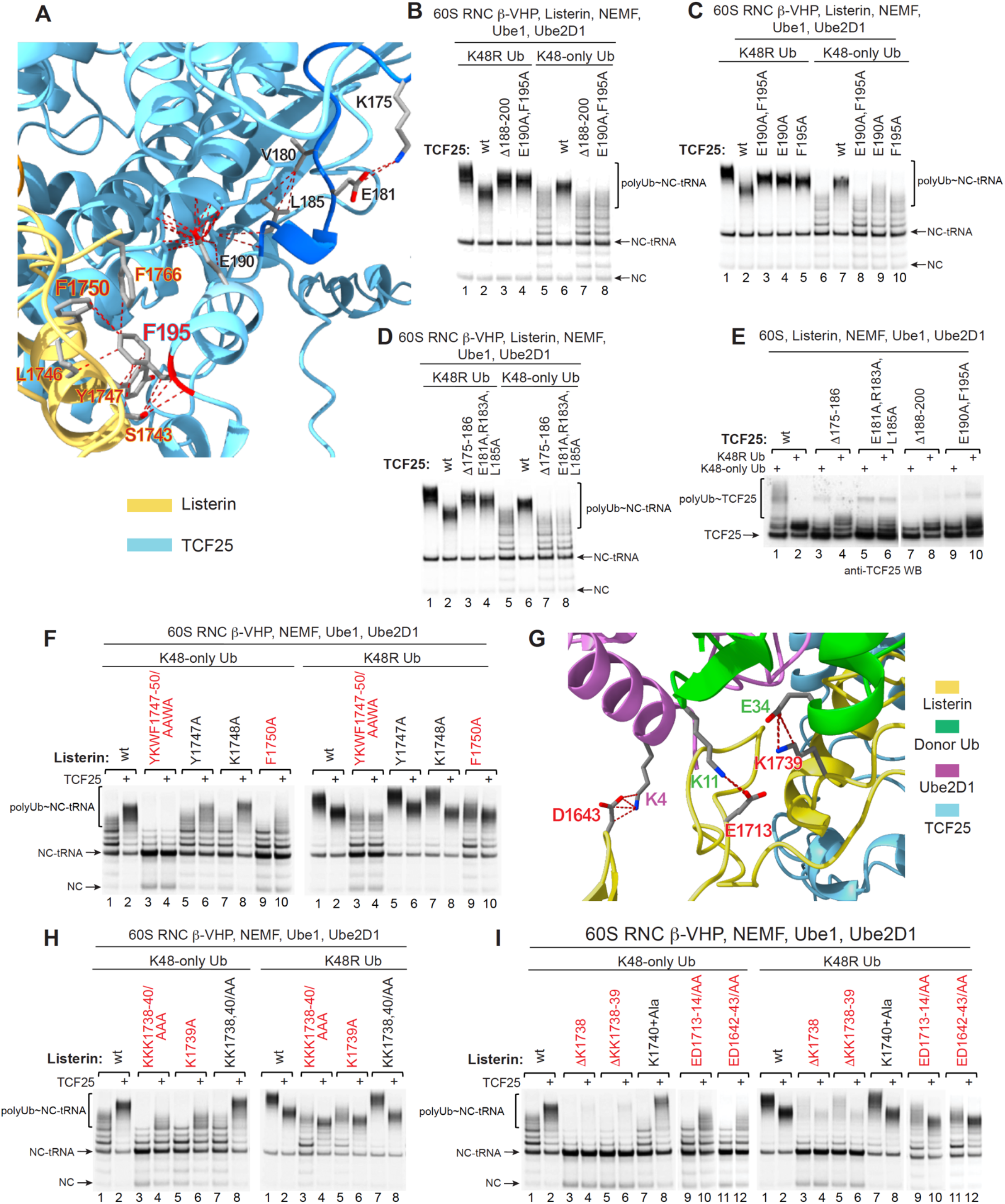
Mutational analysis of predicted interactions involving Listerin. (A) The area of the predicted interaction between TCF25 (light blue ribbon) and the RING domain (yellow ribbon). The potential contacts between specific amino acids (shown in sticks) are rendered as red dotted lines. (B-D) The influence of TCF25 containing mutations that are predicted to affect the TCF25/RING domain interaction (panel A) on ubiquitination of ϕ-VHP-containing 60S RNCs by Listerin/NEMF/Ube1/Ube2D1 with K48-only or K48R mutant Ub, assayed by SDS- PAGE and visualized by Phosphoimager. (E) Listerin-mediated ubiquitination of TCF25 containing mutations that are predicted to affect the TCF25/RING domain interaction (panel A) in reaction mixtures containing NEMF/Ube1/Ube2D1/60S subunits and K48-only or K48R mutant Ub, assayed by western blotting. (F) The activity of Listerin containing mutations that are predicted to affect the TCF25/RING domain interaction (panel A) in ubiquitination of ϕ-VHP-containing 60S RNCs in reaction mixtures with NEMF/Ube1/Ube2D1 and K48-only or K48R mutant Ub in the presence/absence of TCF25, assayed by SDS-PAGE and visualized by Phosphoimager. (G) The areas of the predicted interactions of Listerin (yellow ribbon) with donor ubiquitin (green ribbon) and Ube2D1 (magenta ribbon). The potential contacts between specific amino acids (shown in sticks) are rendered as red dotted lines. (H-I) The activity of Listerin containing mutations that are predicted to affect its interaction with donor ubiquitin and Ube2D1 (panel G) in ubiquitination of ϕ-VHP-containing 60S RNCs in reaction mixtures with NEMF/Ube1/Ube2D1 and K48-only or K48R mutant Ub in the presence/absence of TCF25, assayed by SDS-PAGE and visualized by Phosphoimager.

To assay the role of the corresponding TCF25-interacting residues in the RING domain, we first introduced simultaneous Y1747A/K1748A/F1750A substitutions. The activity of this mutant was very strongly reduced, and importantly, it totally lost the ability to respond to TCF25 (Figure 4F, lanes 3-4 in both panels). Individually, the K1748A and L1746A substitutions did not affect the activity of Listerin or its response to TCF25, whereas the Y1747A substitution had only a small inhibitory effect (Figure 4F, lanes 5-8 in both panels; Figure S3C). In contrast, the ability of the F1750A mutant to respond to TCF25 was strongly reduced (Figure 4F, lanes 9-10 in both panels), underscoring the importance of the F195-F1750 interaction. Other predicted interactions between TCF25 and the RING domain that were not tested include e.g. contacts of K511 and D512 in the loop between helices α23 and α24 in TCF25 with D1763 and R1735 in the RING domain, respectively.

Next, we assayed the importance of several predicted interactions between Listerin and E2 and Ub components of the Ube2D1∼Ub^D^ conjugate. The Listerin RING domain has the unusual coordinating site for the second Zn^2+^ that contains Cys at the fourth position (Cys1/Cys2•His5/Cys6, Cys3/***<u>Cys4</u>***•Cys7/Cys8), forming a PHD finger (Aasland et al., 1995) (Figures S3A-B). Moreover, the loop between Cys4 and His5 is unusually long (4 instead of 2-3 amino acids) and contains conserved Lys residues KKK1738-1740 (Figures S3A-B). In the model, K1739 is predicted to interact with E34 of Ub^D^ (Figure 4G). Simultaneous alanine substitution of all three Lys residues very strongly reduced the activity of Listerin (Figure 4H, lanes 3-4 in both panels). Consistent with the prediction, the K1739A substitution alone had the same strong inhibitory effect (Figure 4F, lanes 5-6 in both panels), whereas the activity of the K1738A/K1740 double mutant was similar to that of the wt Listerin (Figure 4F, lanes 7-8 in both panels). Shortening the Cys4- His5 loop by one or two Lys residues nearly abrogated the activity of Listerin (Figure 4I, lanes 3-6 in both panels), whereas insertion of Ala into this loop after K1740 was well tolerated (Figure 4I, lanes 7-8 in both panels). These results establish the importance of the extended size of this loop and the interaction of its K1739 constituent with Ub^D^ for Listerin’s function. Another predicted interaction between Listerin and Ub^D^ involves E1713 and K11, respectively (Figure 4G). Alanine substitution of the conserved E1713 and D1714 substantially reduced the activity of Listerin (Figure 4I, lanes 9-10 in both panels). We also noticed that Ube2D1 interacts not only with the RING domain but also establishes a contact with the immediately preceding RWD domain through K4 in Ube2D1 and D1643 in this domain (Figure 4G). Alanine substitution of the conserved neighboring ED1642-1643 also reduced the activity of Listerin (Figure 4I, lanes 11-12 in both panels). Importantly, all these Listerin mutants containing substitutions in the regions interacting with Ub^D^ and Ube2D1 were able to respond to TCF25.

Mutation of key molecular linchpin residues required for initiating allosteric activation of the E2∼Ub^D^ conjugate completely inactivates some E3 ligases and strongly reduces the activity of others (Pruneda et al., 2012; Lips et al., 2020; Nakasone et al., 2022). In Listerin, Ala substitution of the corresponding R1762 did not abolish, but very strongly reduced Listerin’s activity (Figure S3E).

According to the model, TCF25 forms multiple contacts with Ub^A^ involving amino acids in several regions in the C-terminal part of TCF25 (Figure 5A, shown in dark blue) as well as in the α9 helix and the loop between helices α7 and α8 (Figure 5A, shown in purple). The model also predicts the interaction of 3_10_-helix Ub^A^ (aa 54-60) with the N-terminal region of α3-helix and a few preceding aa in Ube2D1 (aa 116- 121) (Figure 5A, shown in hot pink). Serial C-terminal deletions (mutants aa 1-633 to aa 1-534) progressively reduced the ability of TCF25 to impose K48-specificity on ubiquitination of itself in reaction mixtures containing 60S subunits, Listerin and NEMF (Figure 5B). Thus, deletion of the very C-terminal binding site (aa 638-642) in the mutants containing aa 1-602 and 1-633 allowed their efficient polyubiquitination by K48R Ub but ubiquitination by K48-only Ub was still stronger (Figure 5B, lanes 3-6), whereas the additional loss of two other C-terminal binding sites in the mutants containing aa 1-551 and 1-534 resulted in their more efficient ubiquitination by K48R than by K48-only Ub (Figure 5B, lanes 7-10). Further truncation (mutants aa 1-511 and aa 1-461) nearly eliminated ubiquitination of TCF25 (Figure 5C). These mutants behaved atypically during purification, which suggests probable misfolding. Consistent with the activity of C-terminally truncated TCF25 mutants in their own ubiquitination, gradual truncation of TCF25 progressively reduced its activity in ubiquitination of nascent chains in 60S RNCs (Figure 5D).

**Figure 5.**
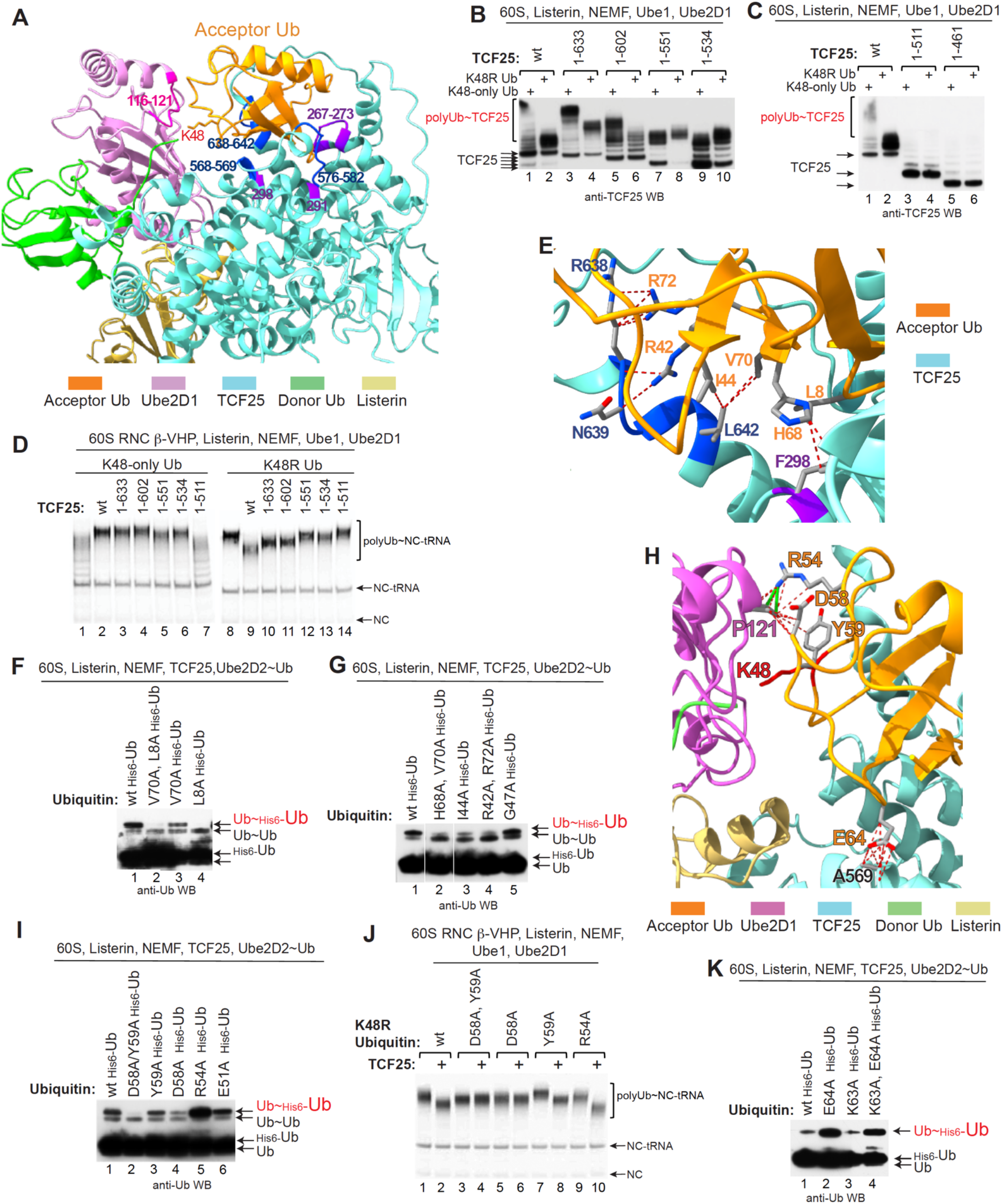
Mutational analysis of predicted interactions involving acceptor ubiquitin. (A) The regions on TCF25 (dark blue and purple on light blue ribbon) and on Ube2D1 (hot pink on pale magenta ribbon) that are predicted to interact with acceptor ubiquitin (orange ribbon). (B-C) Listerin-mediated ubiquitination of C-terminally truncated TCF25 mutants in reaction mixtures containing NEMF/Ube1/Ube2D1/60S subunits and K48-only or K48R mutant Ub, assayed by western blotting. (D) The influence of C-terminally truncated TCF25 mutants on ubiquitination of ϕ-VHP-containing 60S RNCs by Listerin/NEMF/Ube1/Ube2D1 with K48-only or K48R mutant Ub, assayed by SDS-PAGE and visualized by Phosphoimager. (E) The area of predicted interactions between TCF25 (dark blue and purple on light blue ribbon) with acceptor ubiquitin (orange ribbon). The potential contacts between specific amino acids (shown in sticks) are rendered as red dotted lines. (F, G, I, K) The activity of His_6_-tagged Ub mutants in TCF25-mediated formation of di-ubiquitin in reaction mixtures containing Listerin, 60S subunits, NEMF and Ube2D2∼Ub conjugate, assayed by western blotting. Separation of lanes 1/2/3 in panel G by white lines indicates that the juxtaposed parts were from the same gel. (H) The areas of the predicted interactions of acceptor ubiquitin (orange ribbon) with Ube2D1 (magenta ribbon) and TCF25 (light blue ribbon). The predicted clash between R54 (shown in stick) in acceptor Ub and P121 (shown in stick) in Ube2D1 is indicated by green solid lines. The potential contacts between specific amino acids (shown in sticks) are rendered as red dotted lines. (J) Listerin-mediated ubiquitination of ϕ-VHP-containing 60S RNCs in reaction mixtures containing NEMF/Ube1/Ube2D1 and various K48R Ub mutants in the presence/absence of TCF25, assayed by SDS-PAGE and visualized by Phosphoimager.

Next, we evaluated the role of R42, R72, V70 and I44 in Ub^A^, which are predicted to interact with the very C-terminal binding region of TCF25 (aa 638-642) (Figure 5E) deletion of which already substantially reduced the activity of TCF25 (Figure 5B and 5D). For this, we employed a di-Ub formation assay, which allows assessment of the activity of even those Ub^A^ residues which are essential for synthesis of E2∼Ub^D^. V70A, I44A and R42A/R72A substitutions all substantially reduced di-Ub formation (Figure 5F, lane 3; Figure 5G, lanes 3 and 4). H68 is positioned to interact with F298 in TCF25 (Figure 5E) and combining H68A with V70A reduced the activity of Ub further compared to the V70A substitution alone (Figure 5G, lane 2). Notably, the L8A substitution completely abolished synthesis of di-Ub (Figure 5F, lane 4). L8 does not seem to interact directly with TCF25 but could be important for maintaining the Ub hydrophobic patch and the overall structure of Ub.

Interestingly, in the area of contacts between the 3_10_-helix of Ub^A^ (aa 54-60) and the N-terminal region of the α3-helix and a few preceding aa in Ube2D1 (aa 116-121), the model predicts not only interaction of D58 and Y59 with Ube2D1, but also a clash between R54 in Ub^A^ and P121 in Ube2D1 (Figure 5H). The double substitution D58A/Y59A nearly abrogated di-Ub formation (Figure 5I, lane 2). Individually, the D58A substitution strongly decreased synthesis of di-Ub, whereas the Y59A substitution had a low effect (Figure 5I). But strikingly, alleviation of the predicted clash by the R54A substitution strongly increased synthesis of di-Ub (Figure 5I, lane 5). The region of Ub containing R54, D58 and Y59 is not involved in the interaction with Ube1 or Ube2D1, which allowed us to assay these mutants in ubiquitination of 60S RNCs. Consistent with their activities in formation of di-Ub, the D58A substitution almost abrogated the ability of TCF25 to inhibit ubiquitination of nascent chains by K48R Ub, whereas the R54A substitution strongly increased TCF25-mediated inhibition (Figure 5J). We also noticed that on the opposite side of Ub^A^, E64 comes in a close contact with the region of TCF25 containing A569 (Figure 5H). Similarly to R54A substitution, substitution to Ala of E64, but not of the neighboring K63, strongly stimulated synthesis of di-UB (Figure 5K). It is possible that this mutation also alleviates the clash between R54 in Ub^A^ and P121 in Ube2D1.

### The role of the N-terminal region of TCF25

Finally, we assayed the role of the mostly unstructured N-terminal region of TCF25 (aa 1-174), which was not predicted to interact with Listerin or Ub^A^. Deletion of the first 45 or 91 amino acids moderately reduced the ability of TCF25 to stimulate ubiquitination of NCs by the K48-only Ub and to inhibit ubiquitination by K48R Ub (Figure 6A). Substitution of stretches of negatively charged amino acids in this region (aa 30- 33, 35-37 and 68-73) by lysines did not influence the activity of TCF25 (Figure 6B-D), whereas mutation of the conserved Arg residues in the predicted N-terminal α-helix (R4D/R7D/R8D) reduced the activity (Figure 6C). Mutation of the basic stretch residues RKKKKK126-131 of the b-HLH motif to Ala or Asp substantially reduced the activity of TCF25 (Figure 6E), whereas deletion of the adjacent aa 134-145 or of the first helix (aa 147-160) of this motif had only a minor effect (Figure 6F). All TCF25 mutants with impaired activity in ubiquitination of nascent chains were also less efficiently ubiquitinated themselves in reaction mixtures containing 60S subunits, Listerin and NEMF, but nevertheless, like the *wt* TCF25, they were polyubiquitinated with K48-only and only monoubiquitinated with K48R Ub (Figures 6G-H). Thus, despite having lower activity, these mutants retained the ability to impose K48 specificity, which suggests that the N-terminal region of TCF25 does not affect its ability to interact with Ub^A^ but might increase its overall affinity to 60S RNCs, e.g. by interacting with 60S subunits.

**Figure 6.**
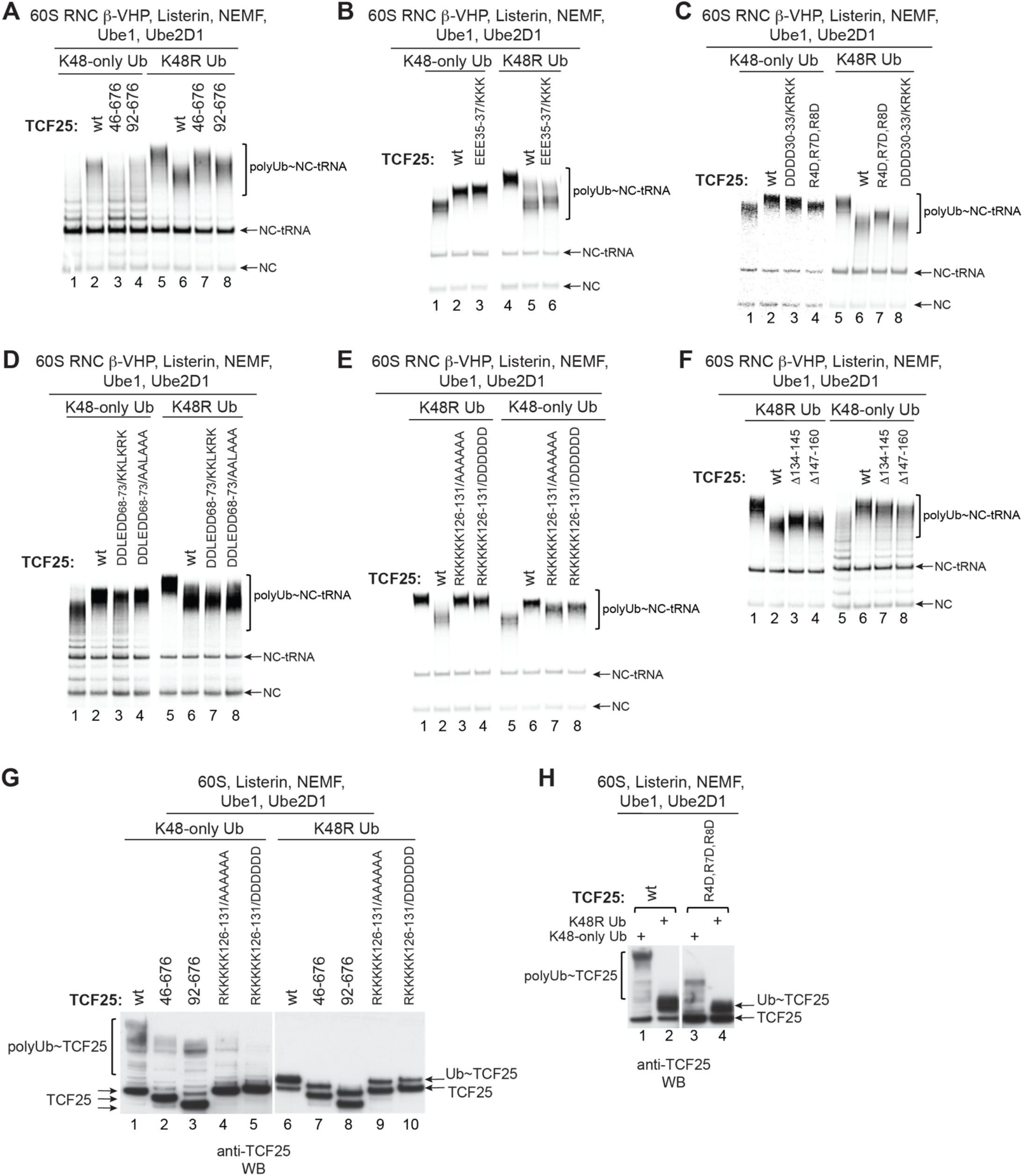
The role of the N-terminal region in the activity of TCF25. (A-F) Influence of *wt* and mutant TCF25 on ubiquitination of ϕ-VHP-containing 60S RNCs by Listerin/NEMF/Ube1/Ube2D1 with K48-only or K48R mutant Ub. Ubiquitination was assayed by SDS-PAGE and visualized by Phosphoimager. Separation of lanes 2/3 and 6/7 in panel D by white lines indicates that the juxtaposed parts were from the same gels. (G-H) Listerin-mediated ubiquitination of *wt* and mutant TCF25 in reaction mixtures containing NEMF/Ube1/Ube2D1/60S subunits and K48-only or K48R mutant Ub, assayed by western blotting.

## DISCUSSION

We have previously determined that TCF25 imposes K48 specificity on ubiquitination by Listerin of nascent chains on 60S RNCs (Kuroha et al, 2018). Here, we present compelling functional data from analysis of the biochemical activities of TCF25 in the *in vitro* reconstituted system and structural evidence (the mutationally tested AlphaFold 3 model of the Listerin/TCF25/Ube2D1/Ub^D^/Ub^A^ complex) that show that TCF25 performs this function by binding Ub^A^ directly and orienting it such that its K48 residue is positioned to attack the thioester bond of the Ube2D1∼Ub^D^ conjugate. We also found that TCF25 itself undergoes K48-specific ubiquitination by Listerin.

Interaction of RING domains with UBC domains of E2 enzymes commonly involve two loop-like regions coordinating Zn^2+^ ions and surrounding a groove formed by the αA helix in the RING domain, and the N-terminal α helix and loops L1 and L2 in the UBC domain (Zheng et al., 2000; Plechanonová et al., 2012; Gundogdu and Walden, 2019). However, additional interactions required for promotion of the closed conformation of the E2∼Ub^D^ conjugate by RING-type E3 ligases are very diverse. In homodimeric RING-type E3 ligases that dimerize by interaction of RING domains, the RING domain of one subunit interacts with E2 enzyme, whereas the RING domain on the second subunit makes contacts with Ub^D^ favoring the closed conformation of E2∼Ub^D^ (Plechanovová et al., 2011, 2012; Dou et al., 2012). In monomeric RING-type E3 ligases, the additional contacts with Ub^D^ that promote closed conformation of the conjugate can be provided by the regions of E3 ligases that are outside of the RING domain, such as a phosphorylated linker tyrosine adjacent to the RING domain of CBL-B (Dou et al., 2013). The monomeric E3 ligase Arkadia is an interesting case in which the activity of the RING domain in promoting the closed conformation of the E2∼Ub conjugate is stimulated by binding of a second, regulatory ubiquitin molecule (Ub^R^) to the surface of the RING domain opposite the E2-binding site: in the resulting quaternary complex, the closed conformation is stabilized by the Ub^D^-Ub^R^ interaction (Wright et al., 2016; Paluda et al., 2022). In the Listerin/TCF25/Ube2D1/Ub^D^/Ub^A^ model, Ube2D1∼Ub^D^ is in the closed conformation, and the interaction between the RING domain and Ube2D1 involves the canonical surfaces. As predicted by the model and supported by our mutagenesis results, the closed conformation of the Ube2D1∼Ub^D^ is strongly stabilized by the interaction of K1739 located in the unusually long Cys4-His5 loop of the RING domain with E34 in Ub^D^ (Figure 5). The presence of Lys at the 1739 position and the extended size of the loop are both essential for Listerin’s function. An additional predicted and tested interaction of Listerin with Ub^D^ involves the contact between E1713 at the N-terminus of the RING domain and K11 in Ub^D^. The interactions between Leu1760 in the RING domain with Leu8 and Ile36 in Ub^D^ are also possible.

According to the model, TCF25 interacts with the RING domain but not with Ube2D1, and the only predicted contact between TCF25 and Ub^D^ is the hydrogen bond between K511 in TCF25 and E39 in Ub^D^, which is unlikely to be sufficient to explain stimulation by TCF25 of Listerin-mediated hydrolysis of Ube2D1∼Ub^D^ by stabilization of its closed conformation. TCF25 binds to the surface of the RING domain opposite the Ube2D1-binding site, and the main interactions include hydrophobic contacts between amino acids in the C-terminal part of the αA helix of the RING domain and in α5-α6 helices of TCF25, with F1750 in the RING domain and F195 in TCF25 being particularly important. The stimulation of Ube2D1∼Ub^D^ hydrolysis by TCF25 could therefore involve its potential influence on the conformation of the RING domain. Stimulation of hydrolysis by enhancement of ribosomal binding of Listerin by TCF25 can also not be excluded.

The main function of TCF25 in Listerin-mediated ubiquitination of nascent chains in 60S RNCs is to impose K48 specificity on ubiquitination, which it performs by binding and orienting the acceptor ubiquitin so that K48 is positioned to attack the thioester bond of the Ube2D1∼Ub^D^ conjugate. Specific essential interaction of TCF25 with the RING domain of Listerin rather than with E2 explains why TCF25 does not perform its function when Ube2D E2s are combined with other E3 ligases (Kuroha et al., 2018). TCF25 forms multiple contacts with Ub^A^ involving amino acids in several regions in the C-terminal part of TCF25, and consistently, progressive C-terminal deletions gradually reduced the ability of TCF25 to impose K48-specificity on ubiquitination of nascent chains and on itself. However, the overall efficiency of ubiquitination of C-terminally truncated TCF25 mutants themselves was not reduced indicating that these deletions did not interfere with TCF25’s binding to the RQC complexes and that their effect was exclusively due to the loss of its specific interaction with Ub^A^. According to the model, the contact area of Ub^A^ with Ube2D1 involves the 3_10_-helix of Ub^A^ (aa 54-60) and the N-terminal region of the α3-helix and a few preceding aa in Ube2D1 (aa 116-121). The model also predicted a clash in this area between R54 in Ub^A^ and P121 in Ube2D1. Strikingly, whereas D58A substitution in Ub^A^ expectedly strongly inhibited the activity of TCF25, consistent with the predicted clash between R54 and P121, its alleviation by the R54A substitution strongly enhanced TCF25’s ability to impose K48 specificity. The general importance of the interaction between the 3_10_-helix in Ub^A^ and the N-terminal region of the α3-helix of the UBC domain for K48 specificity of ubiquitination has been reported previously for other E2 enzymes, e.g. Ube2K in which the position of Ub^A^ is stabilized not by a separate protein, but by Ube2K’s C-terminal UBA domain (Middleton and Day, 2015; Nakasone et al., 2022).

Our finding that TCF25 itself can undergo K48-specific ubiquitination by Listerin yielding proteasome-degradable polyubiquitinated TCF25 suggests a mechanism for the upregulation of yeast Rqc1 levels in the absence of Ltn1 (Barros et al., 2021) and the observed degradation of mammalian TCF25 by the proteasome *in vivo* (Dos Santos et al., 2024). The fact that mutations of TCF25’s ubiquitination sites did not influence its activity in ubiquitination of nascent chain indicates that Listerin-mediated ubiquitination of TCF25 acts only as the instrument to control cellular levels of TCF25. Most ubiquitination sites in TCF25, except K231, occur in its unstructured (likely flexible) N-terminal regions that are not predicted to be involved in interactions with the RING domain or Ub^A^. It is therefore likely that they can be ubiquitinated on the TCF25 molecule that is accommodated in the RQC complex. However, K231 is located on the structured part of TCF25 far from the thioester bond of the Ube2D1-Ub^D^ conjugate and would probably have to be ubiquitinated before full accommodation of TCF25. The fact that some N-terminal regions that were not predicted to interact with the RING domain or Ub^A^ can influence the overall efficiency of TCF25 indicates that they could be involved in interactions that increase TCF25’s affinity to RQC complexes, e.g. by interacting with 60S subunits. Such interactions without full accommodation of TCF25 might also allow ubiquitination of K231.

In the absence of 60S subunits and NEMF/TCF25, Listerin did not stimulate hydrolysis of the Ube2D1-Ub conjugate, indicating that it existed in an in auto-inhibited conformation, which is common for E3 ligases. However, we sometimes purified Listerin* that could undergo self-ubiquitination in the presence of only Ube1, Ube2D1 and Ub. Moreover, auto-ubiquitination of normal Listerin could be induced by some local unfolding after addition to reaction mixtures of the anionic detergent Sarkosyl at low concentrations. Self-ubiquitination of yeast Ltn1 *in vivo* that reduced cellular Ltn1 levels has also been reported, and rapid proteasome-dependent turnover of Ltn1 occurred not only upon starvation but also in rich medium (Ossaher-Nazari et al., 2014). We could not unambiguously identify circumstances that led to the appearance of Listerin* but noticed a possible correlation with the duration of cell incubation. Thus, ubiquitination of the RQC components *in vivo* and its influence on their levels merits further investigation.

The AlphaFold3 model of the Listerin/TCF25/Ube2D1/Ub^D^/Ub^A^ complex was generated without its naive context, i.e. association with 60S ribosomal subunits. In yeast, the RING domain of Ltn1 was observed in two extreme positions, either close to the 60S (IN) or far away from it (OUT) (Tesina et al., 2023). The AlphaFold3 model is generally compatible with the OUT position of the RING domain, but future studies should concentrate on visualization of the Listerin/TCF25/Ube2D1/Ub^D^/Ub^A^ complex on the 60S subunits. It would be also interesting to determine whether some elements of the N-terminal region of TCF25 could form stable interactions with the components of the complete RQC complex.

## MATERIALS AND METHODS

### Plasmids

The ϕ-VHP transcription vector has been described (Shao et al., 2013). Analogous variants with Lysine codons substituted by CGG (Arginine) codons and novel Lysine codons inserted at positions 30 to 54 from the C-terminus were made by Genscript (Piscataway, NJ). β-VHP plasmids were linearized with SalI to transcribe mRNA ending 2nt after the last sense codon. mRNAs were transcribed using T7 RNA polymerase.

Vectors for mammalian expression of 3XFLAG-tagged *wt* human Listerin, NEMF and TCF25 have been described (Shao and Hegde, 2014; Shao et al., 2015; Kuroha et al., 2018). pcDNA3.1-based vectors for mammalian expression of 3XFLAG-tagged Listerin and TCF25 mutants were made by Genscript. Vectors for expression in *E. coli* of human N-terminally His_6_-tagged Ube2D1 and Ube2D2, and variants thereof were made by Genscript by inserting appropriate DNA sequences between NdeI and BamHI sites of pET3a. Vectors for expression in *E. coli* of N-terminally His_6_-tagged human ubiquitin and mutants thereof were made by inserting appropriate DNA sequences between NdeI and BamHI sites of pET3a (Synbio, Monmouth Junction, NJ).

### Purification of Listerin, NEMF, TCF25, Ube2D1/2, ubiquitin and 60S subunits

Expi293 cells (ThermoFisher Scientific) or HEK293 cells (ATCC) were transfected with vectors for expression of *wt* and mutant forms of Listerin, NEMF and TCF25. HEK293T cells were cultured in Dulbecco’s Modified Eagle Medium (GIBCO), supplemented with 10% FBS (GIBCO), 1X Glutamax (GIBCO) and 1x penicillin/streptomycin (GIBCO) at 37°C with 5% CO_2_. Expi293 cells expressing (a) TCF25 and (b) NEMF and Listerin were grown for 72 and 96 hours, respectively in Expi expression medium (ThermoFisher Scientific).

Cells expressing NEMF or TCF25 were lysed in buffer A (20 mM Tris-HCl pH7.5, 200 mM KCl, 2.5 mM MgCl_2_, 2 mM DTT) supplemented with cOmplete EDTA-free protease inhibitor cocktail (Roche) and 1% Triton X100. The supernatant obtained after centrifugation of lysate for 20 min at 10,000 rpm at 4°C was loaded onto anti-DYKDDDDK G1 affinity resin (GenScript) equilibrated with buffer A. Protein-bound resins were washed sequentially with buffer A, with buffer B (20 mM Tris-HCl pH7.5, 600 mM KCl, 2.5 mM MgCl_2_, 2 mM DTT) and again with buffer A + 15% glycerol (for TCF25) or 10% glycerol (for NEMF). Proteins were eluted by incubation with 0.5 mg/ml 3XFLAG peptide (ApexBio, Houston TX) in buffer A for 1h at 4°C, followed by centrifugation.

Cells expressing Listerin were lysed by Dounce homogenization in hypotonic buffer (20 mM Tris- HCL pH 7.5, 10 mM KCl, 1.5 mM MgCl_2_, 2.5 mM DTT, 0.1 mM EDTA) supplemented with Halt™ Protease and Phosphatase Inhibitor Cocktail (100X) (Thermo Scientific). The KCl content was then adjusted by addition of similarly supplemented hypertonic buffer (20 mM Tris-HCl pH 7.5, 1 M KCl, 1.5 mM MgCl_2_, 2.5 mM DTT, 0.1 mM EDTA). The supernatant obtained after centrifugation of lysate for 30 min at 45,000 rpm in a Beckman SW50.2Ti rotor at 4°C was loaded onto anti-DYKDDDDK G1 affinity resin equilibrated with buffer A. Protein-bound resins were washed sequentially with buffer A, with buffer B and again with buffer A + 10% glycerol. Proteins were eluted by incubation with 0.5mg/ml 3XFLAG peptide in buffer A for 1h at 4°C, followed by centrifugation.

Ube2D1 and Ube2D2 were expressed in 2L *E. coli* BL21(DE3). Protein expression was induced with 1 mM IPTG, and cells were grown for 4 h at 20°C. The cell pellet was resuspended in buffer C (20 mM Tris-HCl pH 7.5, 10% w/v glycerol, 200 mM KCl) supplemented with 2 mM AEBSF (4- benzenesulfonyl fluoride hydrochloride) protease inhibitor. All proteins were purified by Ni-NTA chromatography followed by FPLC on a MonoS 5/50 GL column in buffer D (20 mM HEPES pH 7.5, 10% w/v glycerol, 2 mM DTT) in a linear 50-500 mM KCl gradient. Ube2D1 and Ube2D2 eluted at 140-165mM KCl.

N-terminally His_6_-tagged ubiquitin mutants were expressed in *E. coli* BL21 (DE3). Protein expression was induced with 1 mM IPTG, and cells were cultivated for 3h at 37°C. Ubiquitin proteins were purified by Ni-NTA affinity chromatography as described for Ube2D1/Ube2D2 and dialyzed overnight against buffer E (20 mM Tris-HCl pH 8, 100 mM KCl, 10% w/v glycerol, 2 mM DTT). 60S ribosomal subunits were purified from rabbit reticulocyte lysate (Green Hectares, Oregon WI) as described (Pisarev et al., 2007).

### Preparation of 60S RNCs

60S RNCs were prepared using a modified version of a previously described protocol (Kuroha et al., 2018). Briefly, 250 nM of different β-VHP mRNA variants were translated in Flexi RRL (Promega) supplemented with 3 μM NEMF in the presence of [^35^S]-methionine for 50 min at 32°C. Reaction mixtures were then subjected to centrifugation in a Beckman SW55 rotor at 53,000 rpm for 95 min at 4°C in 10%–30% linear sucrose density gradients prepared in buffer F (20 mM Tris-HCl pH 7.5, 100 mM KCl, 1.5 mM MgCl_2_, 2 mM DTT, 0.25 mM spermidine).

### Ube2D1/2∼Ub conjugates

Ube2D1^C85S^∼Ub oxyester conjugate was prepared as described (Wright et al., 2016). *Wt* Ube2D2∼Ub thioester conjugate was purchased from Bio-Techne (Minneapolis, MN).

### Ubiquitination of 60S RNCs

Nascent chain peptides were ubiquitinated essentially as described (Shao et al., 2013, Kuroha et al., 2018). 5 nM SDG-purified [^35^S]Met-labeled 60S RNCs, 60 nM *wt* or mutant Listerin, 60 nM NEMF, and 60 nM *wt* or mutant TCF25 were preincubated in buffer F supplemented with 1 mM ATP, 1 mM GTP, 2 mM MgCl_2_ to compensate for NTPs, 12 mM creatine phosphate, 2 U/μl RiboLock RNase inhibitor, on ice for 15 min. Reaction mixtures were then supplemented with the indicated combinations of 60 nM Ube1, 240 nM Ube2D1, 1.45 μM ubiquitin (K48R or K48-only), 15.22 μg/ml creatine kinase and incubated for 5 min at 30°C. In all experiments, 20 μl samples were resolved by 4−12% SDS−PAGE, and visualized by Phosphorimager.

### Ube2D1^C85S^∼Ub oxyester conjugate hydrolysis assay

The Ube2D1^C85S^∼Ub oxyester conjugate hydrolysis assay was adapted from a published protocol (Wright et al., 2016). 200 nM of Listerin, 60S subunits, TCF25 and NEMF in different combinations were preincubated on ice for 15 min, after which reaction mixtures were supplemented with 100 nM Ube2D1^C85S^∼Ub conjugate and incubated at 37°C for the indicated times. Reactions were stopped by addition of loading buffer. Samples were resolved by 4-12% SDS-PAGE and visualized by immunoblotting using antibodies against ubiquitin.

### Listerin auto-ubiquitination assay

100-200 nM of Listerin, 5.8 µM *wt* or mutant ubiquitin, 330 nM Ube1, 3.75 µM of Ube2D1 or other E2 enzymes (Bio-Techne; LifeSensors, Malvern, PA) (as indicated in Figures 1E and S2B), 60.87 µg/ml creatine kinase were mixed in buffer F supplemented with 1 mM ATP, 1 mM GTP, 2 mM MgCl_2_ to compensate for NTPs and 12 mM creatine phosphate and incubated for 10 min at 37°C. Some reaction mixtures (Figures S2B and 1G) were additionally supplemented with 600 nM TCF25, NEMF, Listerin or different concentrations of Sarkosyl, as indicated. The reaction was stopped by adding loading buffer. Samples were resolved by 4-12% SDS-PAGE and visualized by Simply blue staining or by western blotting using antibodies against Listerin.

### Ubiquitination and proteasomal degradation of TCF25 *in vitro*

Ubiquitination of *wt* and mutant TCF25 by Listerin was done using vacant 60S subunits. Different combinations of 400 nM Listerin, 400 nM NEMF, 800 nM 60S subunits and 300 nM TCF25 were preincubated on ice for 15 min in buffer F supplemented with 1 mM ATP, 1 mM GTP, 2 mM MgCl_2_ to compensate for NTPs and 12 mM creatine phosphate. Then reaction mixtures were supplemented with 23.2 µM wt, K48R or K48-only ubiquitin, 400 nM Ube1, 3.7 5µM Ube2D1, and 34.78 µg/ml creatine kinase and incubated at 37°C for 10 min. Reactions were stopped by adding loading buffer and divided into two prior to 4-12% SDS-PAGE. Ubiquitination was visualized in parallel by Simply blue staining and by western blotting using antibodies against TCF25.

For the proteasomal degradation assay, after ubiquitination, reaction mixtures were directly supplemented with 40 nM 26S proteasome (Bio-Techne) and incubated at 37°C for another 20 min. Samples were resolved by 4-12% SDS-PAGE and visualized by western blotting using antibodies against TCF25.

### FLAG-tag pull down assays

Control Expi293 cells and cells transiently expressing FLAG-tagged TCF25 were collected, washed with PBS, subjected to Dounce homogenization, and the KCl content of lysates was then adjusted to 100mM KCl. The supernatant obtained after centrifugation of lysate for 10 min at 10,000 rpm at 4°C was loaded onto anti-DYKDDDDK G1 affinity resin equilibrated with buffer G (20 mM Tris-HCl pH7.5, 100 mM KCl, 2.5 mM MgCl_2_, 2 mM DTT). Columns were washed with 10 ml buffer G, and 10 ml buffer G containing 0.5 mg/ml recombinant ubiquitin and 0.5 mg/ml BSA was applied to each column 4 times. The resin was washed again and then resuspended in an equal volume of buffer G. Loading buffer was added to aliquots of resin suspension and samples were resolved by 4-12% SDS-PAGE and visualized by immunoblotting with antibodies against ubiquitin.

### Di-ubiquitin formation assay

200 nM Listerin, 200 nM NEMF, 200 nM 60S subunits, 200 nM TCF25 and 11.6 µM of indicated His_6_- tagged ubiquitin mutant were incubated in buffer F on ice for 15 min. 2.5 µM Ube2D2∼Ub thioester conjugate was then added, and reaction mixtures were incubated at 37°C for 10 min. The reaction was stopped by adding loading buffer. Samples were resolved by 4-12% SDS-PAGE and visualized by immunoblotting using antibodies against ubiquitin.

### AlphaFold3 modeling

The model for the interaction of full-length human TCF25, Listerin, Ube2D1 and two ubiquitin molecules was generated using AlphaFold3 (Abramson et al., 2024).

## COMPETING INTEREST STATEMENT

The authors declare no competing interests.

## ACKNOWLEDGMENTS

We thank Manu Hegde and Susan Shao for pcDNA-based vectors for expression of NEMF and Listerin and protocols for purification of these proteins, and to Andrew Tcherepanov for expert technical assistance. This work was supported by NIH grant GM122602 to T.V.P, and NIH grant GM097014 to C.U.T.H.

## AUTHOR CONTRIBUTIONS

I.S.A. and A.G.B. performed all experiments. T.V.P., C.U.T.H., I.S.A., and A.G.M. designed experiments and interpreted data. T.V.P. wrote the paper with input from all authors.

